# The Celiac Microbiome Repository (CMR): A Curated Collection of Celiac Disease Gut Microbiome Sequencing Data

**DOI:** 10.64898/2026.03.28.715053

**Authors:** Haig V. Bishop, Peter J. Prendergast, Craig W. Herbold, Renwick C.J. Dobson, Olivia J. Ogilvie

## Abstract

Celiac disease is an autoimmune condition where the gut microbiome is increasingly recognised as a key environmental factor. While high-throughput sequencing has led to a surge in celiac-related gut microbiome profiling data, these datasets remain fragmented, heterogeneous, and often lack the metadata required for large-scale integration into pooled, cross-cohort datasets. To address this, we developed the Celiac Microbiome Repository (CMR), a curated, open-access collection of celiac-related 16S rRNA gene and shotgun metagenomic sequencing datasets. We employed a systematic curation workflow to identify datasets across the NCBI Sequence Read Archive (SRA) and Scopus, followed by manual metadata extraction and direct author engagement. All 16S data was reprocessed through DADA2 and shotgun data through MetaPhlAn4 to facilitate comparison across studies. The CMR version 1.0 comprises 28 datasets containing 3,245 samples from 13 countries and 5 body sites. Our analysis reveals that while publicly available celiac microbiome samples have accumulated at a rate of approximately 140 per year, significant barriers to accessibility exist. Just 20 of 58 eligible datasets were found to have both raw data and essential metadata readily available within public archives. The repository features a dual-interface design, consisting of a GitHub backend for programmatic access and an R Shiny frontend for interactive data exploration. By providing this curated and harmonised resource, the CMR enables the research community to leverage public data for global meta-analyses and machine learning applications. Ultimately, this work provides the foundation needed to move beyond isolated, small-scale studies toward high-powered discoveries in celiac disease research.

**Database URLs:** https://github.com/CeliacMicrobiomeRepo/celiac-repository | https://celiac.shinyapps.io/celiac-webapp

## Introduction

Celiac disease is a chronic autoimmune disease of the small intestine triggered by the ingestion of gluten [1–3]. While it originates in the gut and is diagnosed via duodenal biopsy [3], the disease manifests as a systemic disorder with wide-reaching clinical impacts [4]. Affecting 1-2% of the global population [5], the condition’s only treatment is a strict, lifelong adherence to a gluten-free diet [6].

The critical genetic element of celiac disease is Human Leukocyte Antigen (HLA)-DQ2 or HLA-DQ8, which occur in 30-40% of the population [7,8]. Although these genetic variants are necessary, they are not sufficient for disease onset. Instead, environmental factors play a role in determining whether an individual goes on to develop a gluten intolerance [3]. While the precise environmental factors are not yet defined, studies increasingly implicate the gut microbiome in celiac disease pathogenesis [9–11]. Consequently, a growing body of research aims to characterise gut microbiome shifts in celiac disease and define their role in pathogenesis.

High-throughput sequencing (HTS) technologies have revolutionised the field of microbiome research, enabling the profiling of bacterial communities through both 16S ribosomal RNA (rRNA) gene amplicon sequencing and shotgun metagenomic sequencing [12]. While 16S rRNA gene sequencing offers cost-effective taxonomic profiling, it is subject to significant technical biases stemming from technical factors including primer choice and variation in 16S rRNA gene copy number across different taxa [13,14]. Shotgun metagenomics overcomes these limitations by capturing the entire genomic content of the microbial community, providing high-resolution taxonomic information as well as a picture of functional potential [15,16].

In celiac research, HTS technology has led to a surge in studies of the gut microbiome composition. To date, as identified in the present study, 58 such investigations have characterised compositional shifts in the microbiome associated with celiac disease development. For instance, Olivares et al. [17] demonstrated that infants who later develop celiac disease exhibit distinct gut microbiota trajectories, characterised by differentiated abundances of *Bifidobacterium* species compared to healthy controls. Bodkhe et al. [18] identify distinct microbial signatures in the duodenum, finding that celiac patients possess a higher abundance of *Megasphaera* and *Helicobacter* compared to healthy controls and first-degree relatives.

When publishing these HTS-based microbiome profiling studies, researchers frequently deposit their raw sequencing data on public databases such as the NCBI Sequence Read Archive (SRA) [19] and the European Nucleotide Archive (ENA) [20]. These repositories have expanded rapidly over the past decade, with SRA reaching 25.6 petabases of data by 2021 [19], and ENA’s raw read dataset collection growing from 1 million in 2016 to over 37 million by 2025 [20]. In principle, this represents a great wealth of data that supports broader integrative meta-analyses and machine learning developments. However, this data is fragmented, inconsistently processed, unorganised, and frequently has incomplete metadata, hindering its exploitation.

Several publicly funded initiatives address this problem by building public databases that better organise microbiome datasets and process them through a standardised pipeline. For instance, the Gut Microbiome Data Repository (GMrepo) is a collection of 890 human gut microbiome sequencing datasets containing in total 118,965 samples [21]. They provide curated metadata for 302 disease phenotypes alongside demographic information such as country, sex and BMI. Other notable examples include the Integrative Human Microbiome Project (iHMP) [22], curatedMetagenomicData [23], MicrobiomeDB [24], MLrepo [25], and most recently, PRIME [26]. Rather than focusing on specific diseases or cohorts, these repositories aggregate gut microbiome data broadly across a diversity of populations. This makes them highly valuable for bridging the gap between raw data availability and practical usage in machine learning applications, reanalyses and meta-analyses.

While these repositories are invaluable, they typically prioritise data volume and programmatically accessible datasets. Consequently, they often provide incomplete coverage of the available data described in the broader literature, making them less applicable to specialised cohorts. Further, specialist metadata, like disease-specific variables, is frequently missing from these databases, remaining buried in original publications or requiring direct contact with the authors for retrieval. These limitations are particularly evident in celiac disease research. Among the platforms mentioned, GMrepo is the most pertinent as it specifically includes the celiac disease phenotype. In contrast, celiac-specific metadata is entirely absent from the other databases, likely due to prioritisation of conditions that offer much higher data volumes like obesity or Type 2 Diabetes. However, with just four studies and a lack of metadata regarding gluten-free diets, GMrepo’s utility for specialised celiac gut microbiome research is limited.

By building specialised repositories for niche cohorts, it is possible to better realise and align with the FAIR principles (findable, accessible, interoperable, and reusable) [27]. These principles mandate that data is discoverable through persistent identifiers, retrievable via standard protocols, consistently formatted to be machine-readable allowing for cross-study integration, and well-documented to ensure its long-term reusability. Adhering to these guidelines is crucial for ensuring that microbiome data remains a useful asset for the research community rather than a dormant archive.

Some projects have been able to achieve this, for example, the SKIOME project enhanced the utility of skin microbiome datasets by providing deeply curated metadata that is often missing from larger databases [28]. Similarly, by systematically searching and reviewing literature to recover raw data and metadata, exhaustive coverage of human respiratory and psychiatric disorder-related microbiome datasets was achieved by ResMicroDb [29] and PsycGM [30]. These initiatives demonstrate that through manual curation, the persistent barriers of fragmentation and metadata gaps can be overcome to create comprehensive gut microbiome data resources for specialised cohorts.

To address these challenges for celiac disease microbiome research and to close the gap between the abundance of raw sequencing data and its practical utility for large-scale analysis, we collected and curated all available datasets into the Celiac Microbiome Repository (CMR). To ensure the repository is comprehensive, we searched both the SRA and Scopus for celiac-related gut microbiome sequencing datasets and when required, we contacted authors directly to request the data. We carefully collected specialised metadata for every sample and processed it through a single standardised pipeline to yield uniform microbiome community profiles. The resulting data was unified into a single, open-access repository in the form of a GitHub repository backend, with an R Shiny web application frontend that allows users to explore the data interactively.

The Celiac Microbiome Repository, therefore, acts as a centralised access point and record of all celiac-related gut microbiome sequencing datasets. It provides consistent microbial community profiles and curated metadata for every sample, enabling the research community to better leverage public data for meta-analysis, machine learning applications and independent exploration.

## Methods

### Data Curation Workflow

Following a systematic data curation pipeline, we populated the CMR with all available celiac-related gut microbiome sequencing datasets (**Fig. 1**). The methodology is structured into four steps: 1) Literature search and eligibility assessment, 2) Raw data acquisition and metadata extraction, 3) Microbial community profiling, and 4) Documentation, version control, and finalisation.

**Figure 1.**
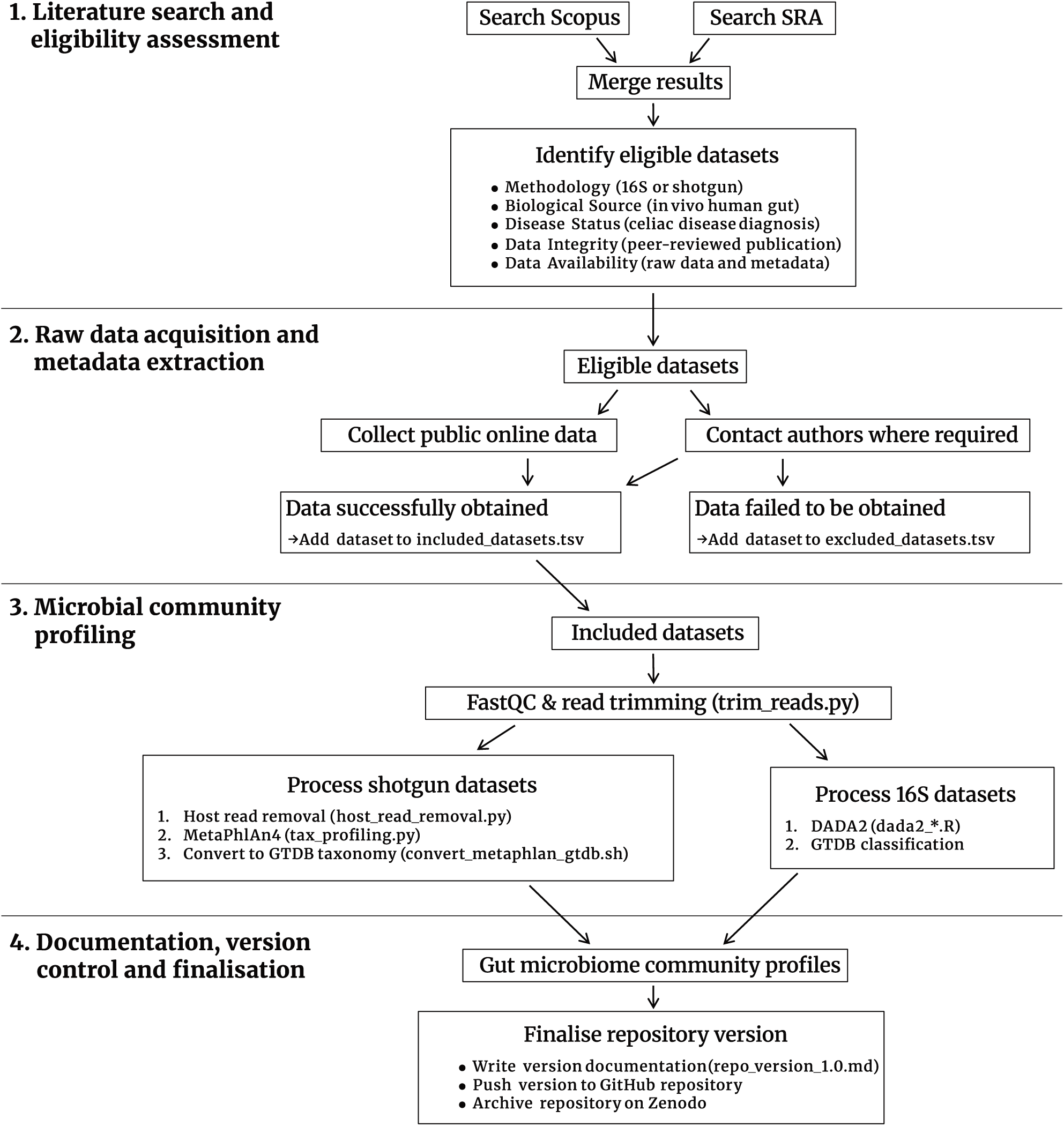
Data curation workflow for the Celiac Microbiome Repository. The four-step pipeline comprises: **1** A dual literature search of Scopus and the NCBI SRA followed by eligibility assessment. **2** Raw data acquisition via public databases and/or direct author contact alongside manual metadata extraction. **3** Microbial community profiling through DADA2 for 16S data and MetaPhlAn4 for shotgun data. **4** Documentation, version control and finalisation including archival on Zenodo. **Alt Text:** Flowchart of the four-step data curation pipeline, proceeding from literature search and eligibility assessment through raw data acquisition and metadata extraction, microbial community profiling, to documentation and archival on Zenodo.

### Step 1: Literature Search and Eligibility Assessment

A systematic literature search was conducted on 15th July 2025. Designed to capture both peer-reviewed publications and raw data deposits, both Scopus and NCBI Sequence Read Archive (SRA) BioProjects were searched. On Scopus, the search was limited to Articles only and used the string: (“celiac”, “coeliac”, “gluten”, “enteropathy” or “sprue”) AND (“metagenomic”, “microbiome”, “microbiota”, “16S”, “rRNA”, “shotgun”, “sequencing” or “metagenomics”). Simultaneously, the SRA was queried for BioProjects using the keywords “celiac”, “coeliac”, “gluten enteropathy”, or “sprue”. These searches aimed to capture all relevant datasets. To be deemed eligible for the CMR, each dataset was evaluated against the following criteria:

#### 1. Methodology

The data must have been generated using either 16S rRNA gene amplicon sequencing or whole metagenomic shotgun sequencing.

#### 2. Biological Source

Only *in vivo* human gastrointestinal microbiome samples are eligible, including, but not limited to, stool, duodenal, gastric, saliva, and oropharynx samples.

#### 3. Disease Status

The dataset must contain samples from individuals who either have already been diagnosed with celiac disease, or longitudinal samples from individuals who later developed the condition (prospective cohorts).

#### 4. Data Integrity

The dataset must be published in a peer-reviewed journal.

#### 5. Data Availability

Either by retrieval from a public database, or by contacting the authors directly, all raw sequencing data and essential metadata must be made available. Essential metadata includes body site, gluten-free diet status and celiac disease status.

SRA records and publications were assigned to one another, and every dataset was carefully inspected to ensure it was indeed eligible for inclusion.

### Step 2: Raw Data Acquisition and Metadata Extraction

The raw sequencing data (per-sample .fastq files), essential metadata (body site, gluten-free diet status and celiac disease status) and other metadata were obtained for each dataset and organised in the GitHub repository according to the CMR’s directory structure (**Fig. 2**). Where available publicly, raw data and metadata were retrieved either from the public database (*e.g.*, the SRA) or from within the publication. The download_sra.py script, a wrapper around the fasterq-dump and prefetch commands from the sra-tools package, automated the retrieval of raw sequencing data from the SRA into a new designated directory.

**Figure 2.**
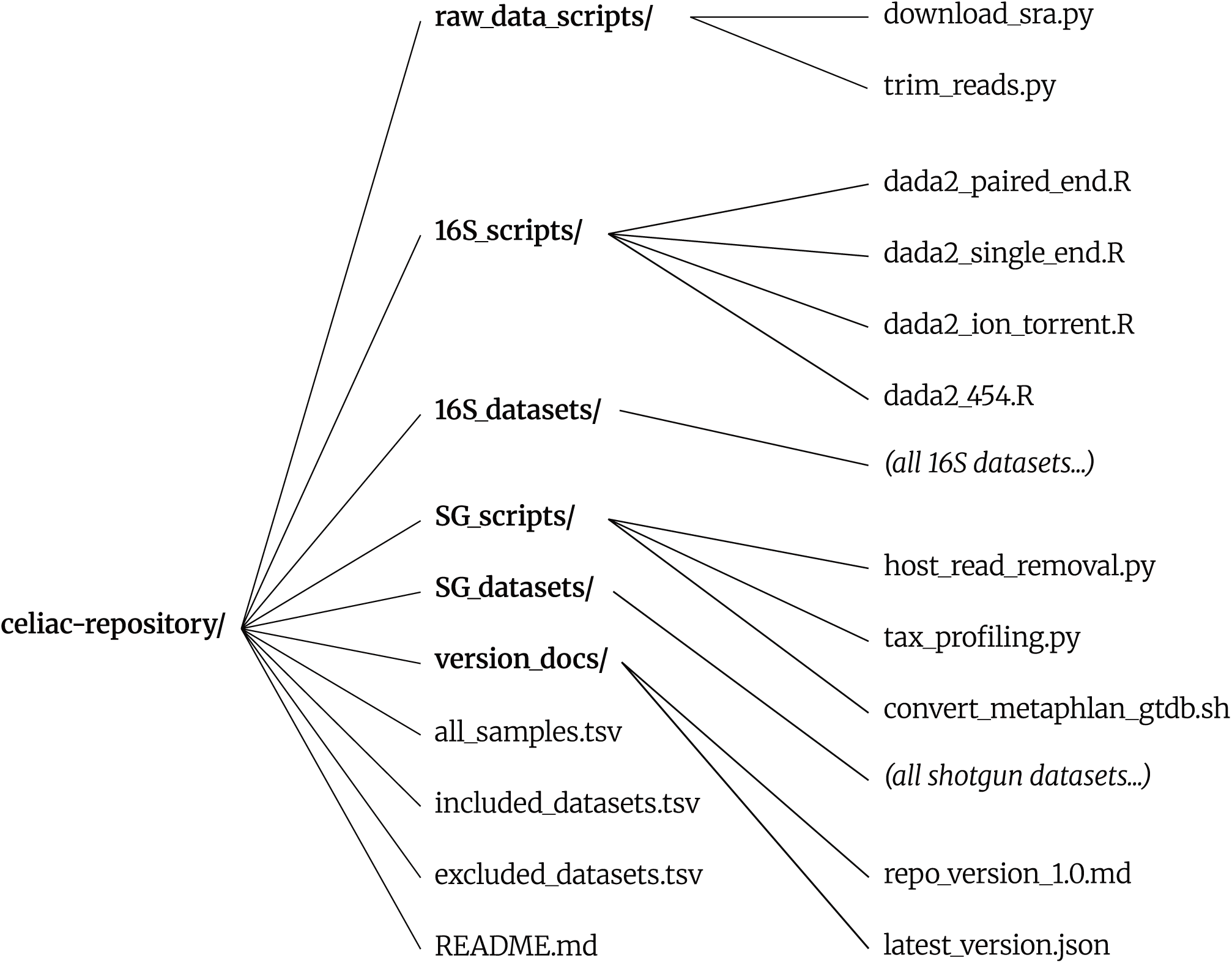
GitHub repository directory structure for the Celiac Microbiome Repository. Python and R scripts for obtaining and processing sequencing data are organised into raw_data_scripts/, 16S_scripts/ and SG_scripts/. All processed sequencing data for every included dataset are stored in 16S_datasets/ and SG_datasets/ according to sequencing methodology. Top-level metadata tables (including included_datasets.tsv, excluded_datasets.tsv, and all_samples.tsv) centralise study and sample metadata. **Alt Text:** Directory tree diagram of the Celiac Microbiome Repository GitHub structure, with subfolders for raw data scripts, 16S scripts, and shotgun scripts, dataset folders organised by sequencing type, and top-level metadata tables at the root.

When not available via the manuscript or a public database, the corresponding author(s) were contacted directly via three successive emails at two-week intervals requesting the raw data and metadata. If no response was received or if the authors were unable to provide the data, the dataset was added to the excluded_datasets.tsv table, along with metadata including publication title, month of publication, Digital Object Identifier (DOI), SRA record (where applicable), sequencing method, reason for exclusion and whether there was a claim of data being available upon reasonable request within the publication.

Successfully obtained datasets were named using a {sequencing_method}_{number_of_samples}_{first_author_last_name} naming convention. The raw data was placed in a new dedicated dataset directory within the repository (*e.g.*, 16S_datasets/16S_24_Bishop/fastqs/) and the included_datasets.tsv and all_samples.tsv tables were updated to contain the metadata for the new datasets and samples. Metadata included a wide range of information about samples and datasets including unique identifiers, publication details, source of the data, country of origin, sequencing methodology (amplicon region, DNA extraction, read pairing, technology, primers, etc.), processing details (populated in Step 3), sample demographics and sample counts, and the essential metadata related to disease status and gluten-free diet. All samples were categorised into six groups according to their disease status, gluten-free diet and study design: active celiac (gluten diet), healthy control (gluten diet), treated celiac (gluten-free diet), treated healthy control (gluten-free diet), and two prospective groups (celiac or healthy) based on future clinical outcomes. Metadata was also recorded regarding significant factors, experimental design-related confounders that were expected to significantly impact the gut microbiome, possibly warranting exclusion or special treatment in downstream analyses. For example, antibiotic usage, non-celiac autoimmune conditions or short-term gluten challenge. A concise study design description was provided for every dataset in a consistent format (*e.g.*, “stool microbiota of TCD with and without other autoimmune conditions”).

### Step 3: Microbial Community Profiling

Handling each dataset separately, the CMR data processing pipeline began with all per-sample raw sequencing .fastq files placed in a directory dedicated to the dataset. Prior to community profiling, the raw sequencing data was quality checked using FastQC. Any adapter sequences were removed using the trim_reads.py script, a wrapper around Trimmomatic and Cutadapt. For shotgun datasets, host read removal was performed using the host_read_removal.py script, a wrapper around Bowtie2. For 16S datasets, DADA2 was used to identify amplicon sequence variants (ASVs) and their relative abundances in the samples. DADA2 recommends different parameters according to the sequencing technology and execution varies according to read pairing. Therefore, 16S datasets were processed using the appropriate script provided by the CMR according to the sequencing technology and read pairing: dada2_454.R for 454 sequencing, dada2_ion_torrent.R for Ion Torrent sequencing, dada2_paired_end.R for paired-end Illumina sequencing, and dada2_single_end.R for single-end Illumina sequencing. For shotgun datasets, the tax_profiling.py script was used to run MetaPhlAn4 to produce taxonomic profiles of the samples. The resulting taxonomic profiles were converted to GTDB (Genome Taxonomy Database) taxonomy using the convert_metaphlan_gtdb.sh script (a wrapper for MetaPhlAn’s sgb_to_gtdb_profile.py script), and both formats were retained in the repository. Processing parameters and options that differed between datasets were recorded in the included_datasets.tsv table, ensuring reproducibility of processing. A summary of all software used in the data curation pipeline and version numbers is provided in **Supplementary Table 1**.

### Step 4: Documentation, Version Control, and Finalisation

To complete the data curation workflow and finalise the CMR update, the following steps were required. The latest_version.json file was updated to hold the correct CMR version number and literature search date. A version markdown document named repo_version_1.0.md was written describing the current version of the CMR in detail and placed in the version_docs/ directory of the repository. The contents of the version document included sections on the literature search results, data acquisition details, high-level overview of the repository, clear definitions for all columns in the metadata tables, and any other relevant information. All .fastq files were removed to avoid their storage in the repository. After local changes were verified, the changes were committed to the Git repository and pushed to the GitHub repository. Upon finalisation of the version, the repository was archived on Zenodo to generate a version-specific DOI, ensuring the data remains findable and citable in its current state.

### ASV Alignment

An in-house alignment tool, extract16s (https://github.com/HaigBishop/extract16s), was used to align ASVs from each dataset to a full-length 16S rRNA gene. Briefly, the barrnap 16S rRNA gene Hidden Markov Model (HMM) was employed to map all ASVs to the coordinates of a single 1525 bp *Escherichia coli* reference 16S rRNA gene (RS_GCF_030545895.1∼NZ_JAUOMX010000042.1:567-2091). To derive a single consensus region for each dataset, low bitscore ASVs and outlier positions were filtered before computing the median start and end positions.

### Analysis of Data Accumulation

To quantify the rate of data accumulation, an Ordinary Least Squares linear regression using the stats.linregress function from SciPy was performed on the cumulative number of celiac disease samples recorded between July 2015 and June 2025 (𝑛 = 120 months). The independent variable was defined as the number of months elapsed from the start of the ten-year window, and the dependent variable was the total cumulative count of available samples from individuals who either have been or will be diagnosed with celiac disease.

## Results

### Dataset Retrieval

The literature search conducted on 15th July 2025, returned 179 results on Sequence Read Archive (SRA) and 927 results on Scopus, ultimately yielding 28 datasets included in version 1.0 of the Celiac Microbiome Repository (**Fig. 3**). Of the 1,106 search results, 58 were confirmed as eligible, 28 of which appeared exclusively on Scopus, while the other 30 appeared on both Scopus and SRA. Using data publicly available online, 24 datasets met the minimal requirements for raw sequencing data and metadata on body site, celiac disease status and gluten-free diet. Four of these datasets required manual collection of metadata from the contents of the associated publication. After contacting authors to request direct access to the missing data, a further four datasets could be added to the repository, bringing the total number of included datasets to 28.

**Figure 3.**
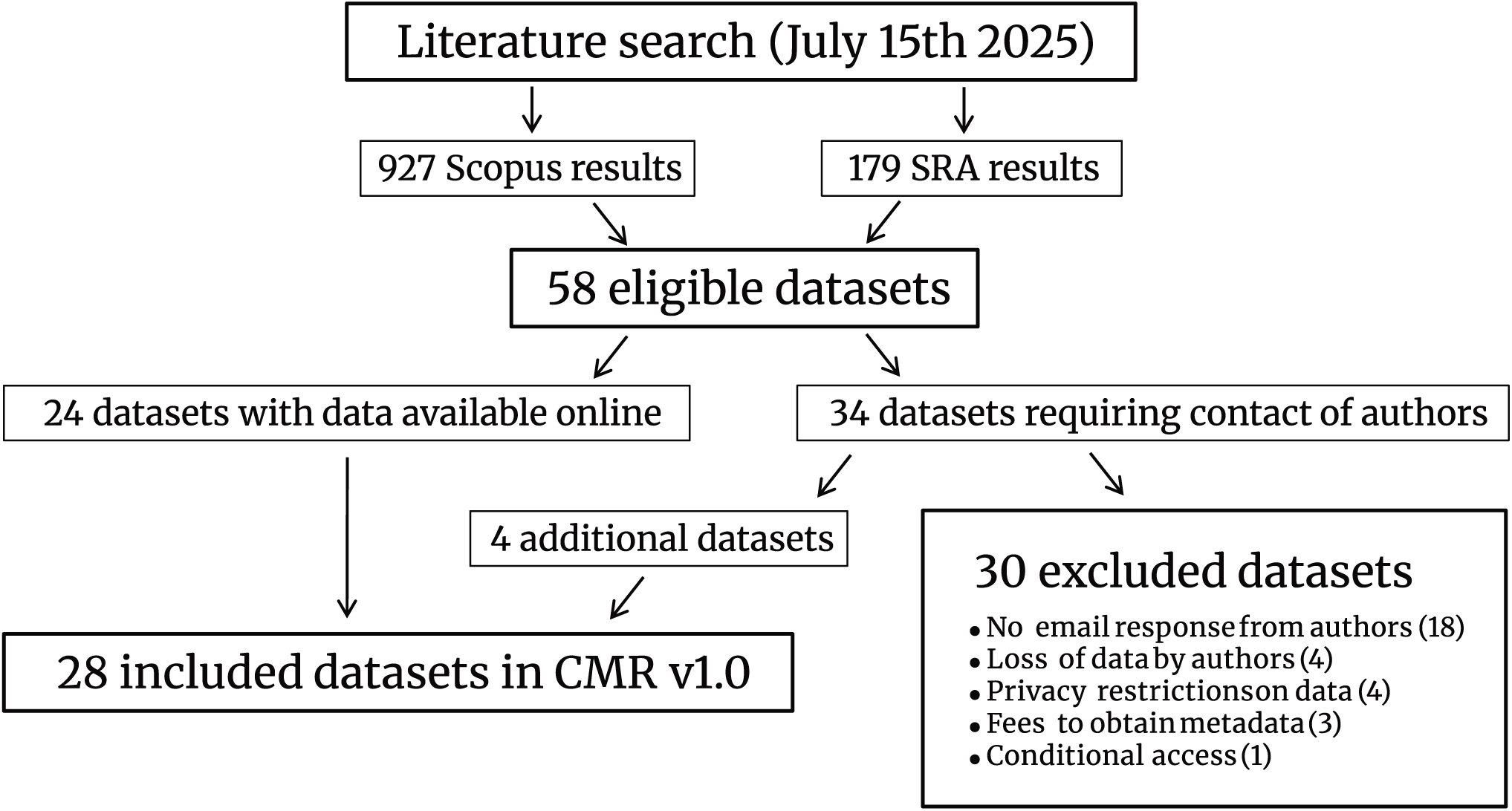
Dataset collection flow diagram. The literature search returned 927 results from Scopus and 179 from the SRA, which after merging yielded 58 eligible datasets. Of these, 24 had data available online and 34 required direct author contact. Author engagement recovered an additional 4 datasets, bringing the total to 28 included datasets. The remaining 30 datasets were excluded due to no email response (𝑛 = 18), loss of data (𝑛 = 4), privacy restrictions (𝑛 = 4), fees to obtain metadata (𝑛 = 3) and conditional access (𝑛 = 1). **Alt Text:** Flow diagram tracking 1,106 combined search results through merging, eligibility screening, and author contact, yielding 28 included datasets. Exclusion reasons shown include no email response, data loss, privacy restrictions, fees, and conditional access.

The remaining 30 datasets (**Supplementary Table 2**) were excluded due to: no email response from authors (𝑛 = 18), loss of data (𝑛 = 4), data privacy restrictions (𝑛 = 4), fees to obtain metadata (𝑛 = 3), and access being conditional on our registration into a formal meta-analysis protocol (𝑛 = 1). Four of the excluded datasets failed to provide data despite claims within the associated publications that it would be made available upon reasonable request.

### Dataset and Sample Overview

The 28 included datasets (**Table 1**) consisted of 3,245 samples spanning 13 different countries and five different body sites (**Fig. 4A-B**). The datasets used a range of body site sampling configurations: stool-only (𝑛 = 17), duodenum-only (𝑛 = 3), stool duodenum (𝑛 = 4), duodenum + stomach (𝑛 = 1), saliva-only (𝑛 = 2), and oropharynx-only (𝑛 = 1).

**Table 1.** Included datasets in version 1.0 of the Celiac Microbiome Repository.

| Dataset ID | BioProject ID | Sequencing Target | Sample Site(s) | Country | Reference |
| --- | --- | --- | --- | --- | --- |
| 16S_136_Nobel | PRJNA778253 | V3–V4 | stool | USA | Nobel et al. [31] |
| 16S_20_Rawson | PRJNA852582 | V3–V4 | stool | USA | Rawson et al. [32] |
| 16S_60_Shi | PRJNA890948 | V3–V4 | stool | China | Shi et al. [33] |
| 16S_96_Quagliariello | PRJEB14943 | V3–V4 | stool | Italy | Quagliariello et al. [34] |
| 16S_27_Fornasaro | NA | V4 | stool | Italy | Fornasaro et al. [35] |
| 16S_49_Turjeman | PRJEB63123 | V4 | stool | Poland | Turjeman et al. [36] |
| 16S_102_Bodkhe | PRJNA385740 | V4 | stool\|duodenum | India | Bodkhe et al. [18] |
| 16S_50_Bibbo | PRJEB35028 | V4 | stool | Italy | Bibbo et al. [37] |
| 16S_227_Oscarsson | PRJNA732664 | V3–V4 | stool | Sweden | Oscarsson et al. [38] |
| 16S_5_Senicar | NA | V3–V4 | stool | Slovenia | Senicar et al. [39] |
| 16S_99_Lionetti | PRJNA923292 | V3–V4 | stool | Italy | Lionetti et al. [40] |
| 16S_80_Garcia | PRJNA401920 | V4 | stool\|duodenum | Mexico | Garcia-Mazcorro et al. [41] |
| 16S_119_Salamon | NA | V3–V4 | stomach\|duodenum | Poland | Salamon et al. [42] |
| 16S_27_Federica | PRJNA750525 | V3–V4 | duodenum | Italy | Ricci et al. [43] |
| 16S_24_Giacomin | PRJNA316208 | V3–V4 | duodenum | Australia | Giacomin et al. [44] |
| 16S_20_Caminero | PRJNA518891 | V3 | duodenum | Canada | Caminero et al. [45] |
| 16S_179_Verdu | PRJNA812940 | V3 | stool\|duodenum | Canada | Constante et al. [46] |
| 16S_25_Francavilla | PRJNA231837 | V1–V3 | saliva | Italy | Francavilla et al. [47] |
| 16S_62_Tian | PRJNA321349 | V3–V4 | saliva | USA | Tian et al. [48] |
| 16S_56_Laffaldano | PRJNA371697 | V4–V6 | oropharynx | Italy | Iaffaldano et al. [49] |
| 16S_39_Olivares | PRJEB23313 | V1–V2 | stool | Spain | Olivares et al. [17] |
| 16S_1211_Milletich | PRJNA854900 | V3–V4 | stool | Sweden | Milletich et al. [50] |
| 16S_68_Girdhar | PRJNA631001 | V4 | stool | Sweden | Girdhar et al. [51] |
| SG_132_Francavilla | PRJNA904924 | Shotgun | stool | Italy | Francavilla et al. [52] |
| SG_80_Mouzan | PRJNA757365 | Shotgun | stool\|duodenum | Saudi Arabia | El Mouzan et al. [53] |
| SG_39_Judith | PRJEB65879 | Shotgun | stool | Spain | Marcos-Zambrano et al. [54] |
| SG_95_Costigan | PRJNA1023737 | Shotgun | stool | England | Costigan et al. [55] |
| SG_118_Leonard | PRJNA486782 | Shotgun | stool | USA\|Italy | Leonard et al. [56] |

All 28 included datasets in version 1.0 of the Celiac Microbiome Repository, where each row reports the dataset identifier, BioProject accession (where available), sequencing target (16S rRNA amplicon region or shotgun metagenomics), sampled body site(s), country, and source publication. NA denotes datasets for which no public BioProject accession was available at the time of curation.

**Figure 4.**
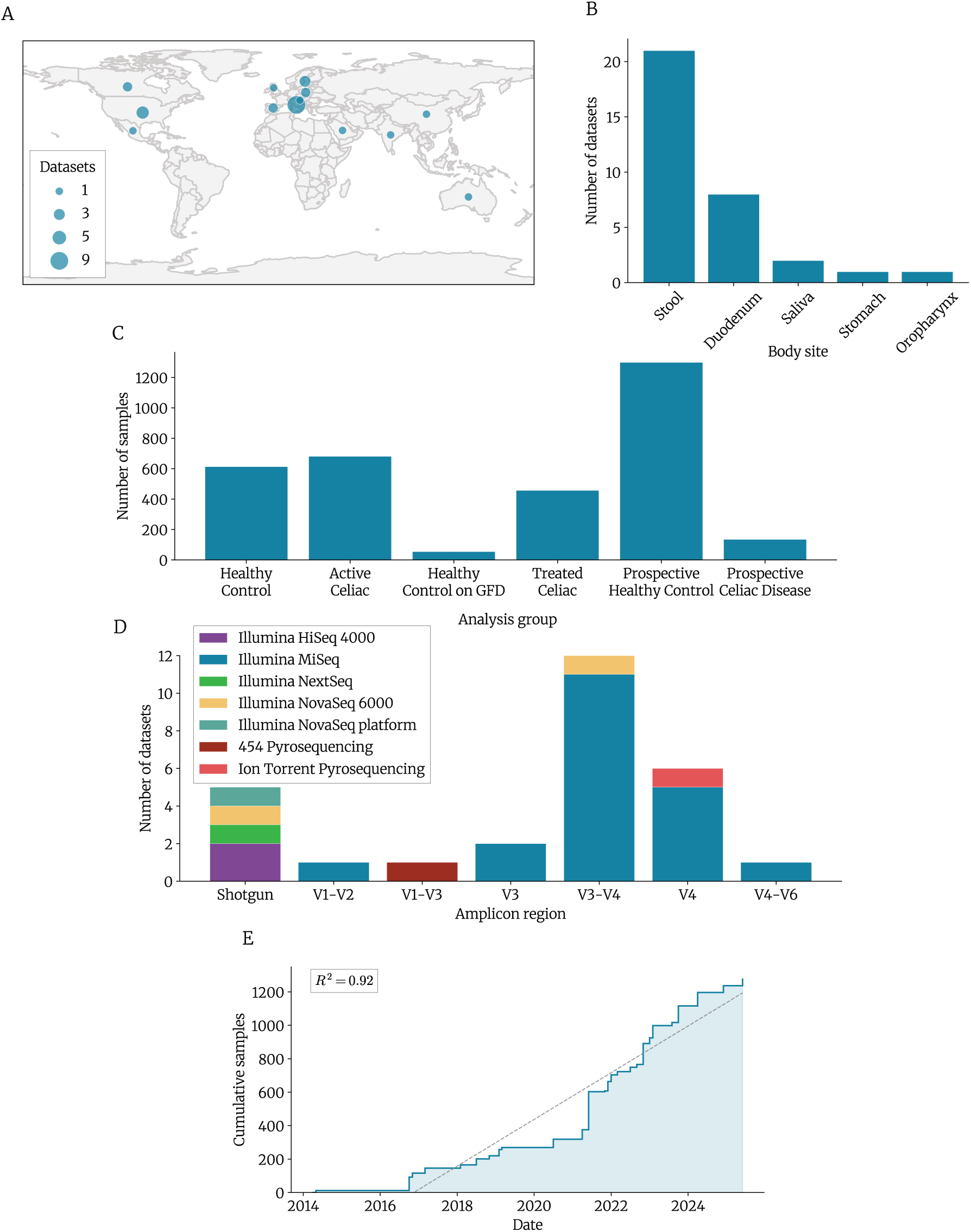
Distributions of the 28 included datasets. **A** Geographic distribution of datasets across countries. **B** Number of datasets by body site. **C** Number of samples per analysis group. **D** Number of datasets by amplicon region and sequencing technology. **E** Cumulative number of publicly available celiac disease samples over the ten-year period from July 2015 to June 2025, with a linear trend line (*R*^2^ = 0.92). **Alt Text:** Five-panel figure showing distributions of the 28 included datasets: a world map of country origins, bar charts of dataset counts by body site and by amplicon region or sequencing technology, a bar chart of samples by analysis group, and a line graph of cumulative publicly available celiac disease samples from July 2015 to June 2025 with a linear trend line.

The majority of the repository consists of cross-sectional data, with 20 datasets utilising 16S rRNA gene amplicon sequencing and four employing shotgun metagenomics. The remaining four datasets follow a prospective design, containing samples from individuals who were followed for several years after sample collection to determine their final celiac disease status. These four datasets comprise three 16S and one shotgun metagenomic dataset, totalling 136 samples from future celiac cases and 1,300 from healthy controls (**Fig. 4C**). For the 24 cross-sectional datasets, all samples could be categorised into four groups according to celiac disease status and gluten-free diet: active celiac (𝑛 = 682), healthy control (𝑛 = 614), treated celiac (𝑛 = 458) and treated healthy control (𝑛 = 55) (**Fig. 4C**).

Dataset-level demographics and significant factor counts are summarised in **Supplementary Table 3**. Five datasets included a total of 157 samples with factors considered as potentially significant biologically relevant confounding factors, including a recent short-term gluten challenge, non-celiac gluten sensitivity, possible celiac disease, a recent hookworm infection and an autoimmune condition other than celiac disease. Sex and age metadata were collected for ten datasets each, including an overlap of eight datasets that contained both sex and age. Datasets that included sex metadata tended to have a higher female-to-male ratio with a mean of 62.3% of samples being female, reflecting the higher prevalence of celiac disease in women [57]. The datasets exhibited a diversity of age ranges, with celiac samples that had available metadata, including 50 infants (0-1 year), 37 children (2-12 years), 7 adolescents (13-17 years), 136 adults (18-64 years), and 10 seniors aged 65 and over.

Five datasets used shotgun metagenomic sequencing, while 23 datasets used 16S rRNA gene amplicon sequencing (**Fig. 4D**). The most common amplicon regions used by the 16S datasets were V3–V4 (𝑛 = 12), V4 (𝑛 = 6) and V3 (𝑛 = 2). Seven different sequencing technologies were used with the most common being Illumina MiSeq, used for 20 16S datasets. Across all datasets, at least 18 unique DNA extraction kits/methods were employed. DNA extraction protocols were classified, revealing that 17 datasets used mechanical lysis in their DNA extraction (*i.e.* bead-beating), ten did not use mechanical lysis and for one dataset the method was not specified.

Based on publication months of included datasets, available celiac disease microbiome samples have accumulated at a rate of 139.64 per year (*R*^2^ = 0.917, 𝑃 < 0.001) over the past ten-year period between July 2015 and June 2025 (**Fig. 4E**).

### Processing Outcomes and Data Quality

After DADA2 processing, median sample read counts across all 16S datasets ranged from 1,943 to 196,162, with an overall median of 29,478 (**Fig. 5A**). Within-sample richness varied across datasets, with median unique ASV counts per sample spanning from 15 to 1,458 (median = 167.5) (**Fig. 5B**). Total dataset-wide richness also showed a broad distribution, with total unique ASV counts falling between 598 and 22,403 (**Fig. 5C**). The alignment of ASVs from all 16S datasets revealed the true 16S rRNA gene regions for each dataset (**Fig. 5D**), where some datasets did not span their entire target 16S rRNA gene hypervariable regions, coinciding with the use of single-end reads.

**Figure 5.**
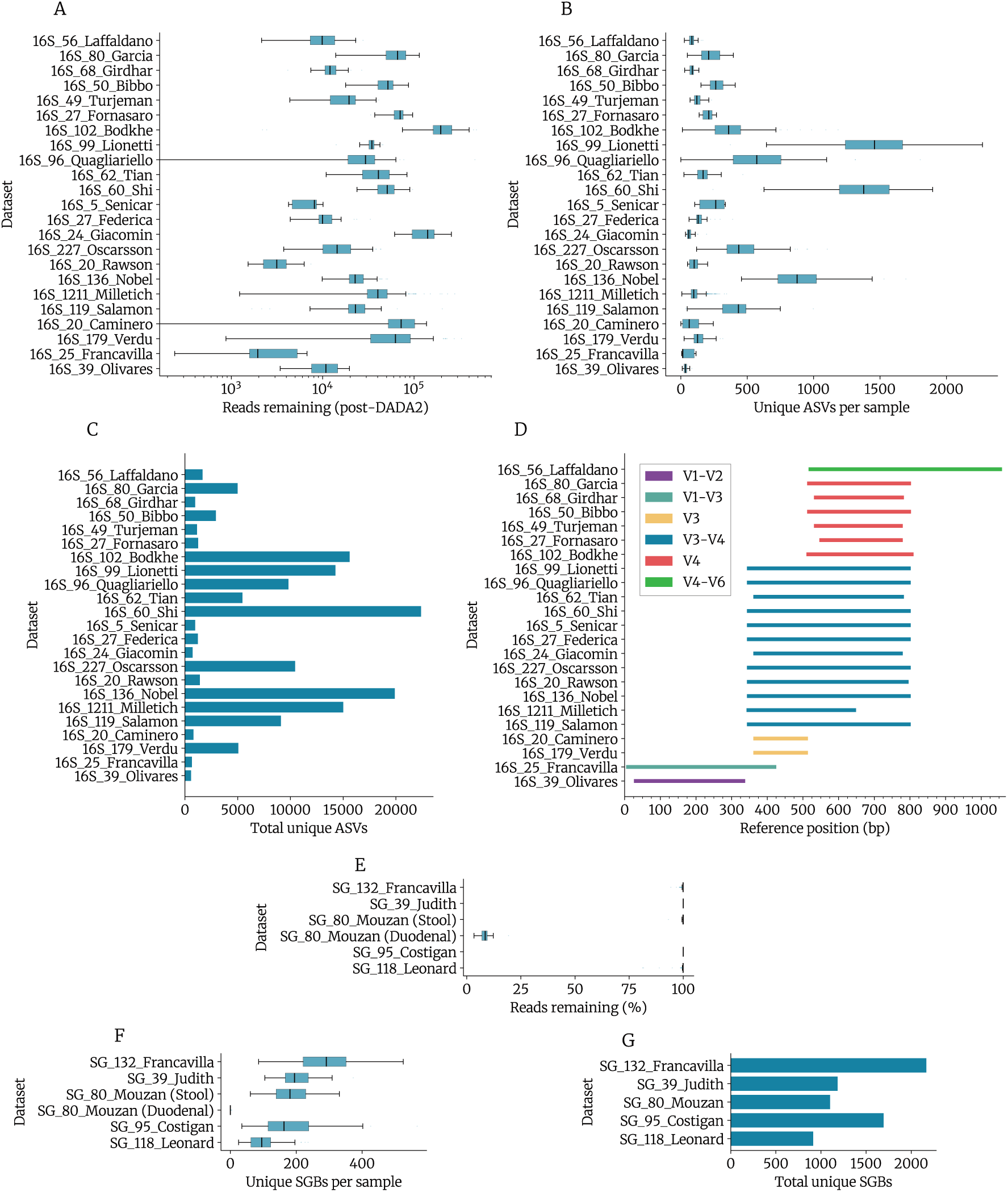
Processing outcomes for all included datasets. **A** Post-processing sample read depth per 16S dataset. **B** Number of unique ASVs per sample per 16S dataset. **C** Total number of unique ASVs per 16S dataset. **D** Alignment of ASVs to a full-length *E. coli* 16S rRNA reference gene, revealing the true amplicon regions per 16S dataset. **E** Percentage of reads remaining after host read removal per shotgun dataset. **F** Number of unique species-level genome bins (SGBs), per sample per shotgun dataset. **G** Total number of unique SGBs per shotgun dataset. **Alt Text:** Seven-panel figure of bioinformatic processing outcomes: post-processing read depth and unique ASV counts per sample and per dataset for 16S data, an alignment plot showing true amplicon regions, host read removal rates for shotgun datasets, and unique species-level genome bin counts per sample and per shotgun dataset.

For shotgun datasets, the percentage of reads remaining after host read removal was almost entirely above 99% for all datasets except for the duodenal samples from SG_80_Mouzan, which ranged between 3.4% and 19.3% with a median of 8.5% reads remaining (**Fig. 5E**). This is explainable by the high host DNA content in the duodenum in combination with shotgun sequencing. The effects of this are also seen in the numbers of unique species-level genome bins (SGB) per sample in each dataset (**Fig. 5F**). The number of unique SGBs per sample for the duodenal samples from SG_80_Mouzan ranged from 0 to 5, while the median results for the remaining datasets ranged between 95.5 and 291.5. Despite these sample-level variations, the total richness across datasets remained high, with the number of unique SGBs ranging from 916 to 2,170 (**Fig. 5G**).

### Repository Interface and Framework

The CMR is primarily implemented as a GitHub repository (https://github.com/CeliacMicrobiomeRepo/celiac-repository). This repository contains all scripts, metadata, processed microbiome profiles, and documentation for the CMR in an open and familiar interface (**Fig. 6A**). To support the CMR GitHub repository, there is also an R Shiny web application that displays the data interactively (https://celiac.shinyapps.io/celiac-webapp). By directly drawing upon the data stored in the GitHub repository, the web application provides a simpler and more user-friendly interface for exploring the data in the form of plots and data tables (**Fig. 6B-C**).

**Figure 6.**
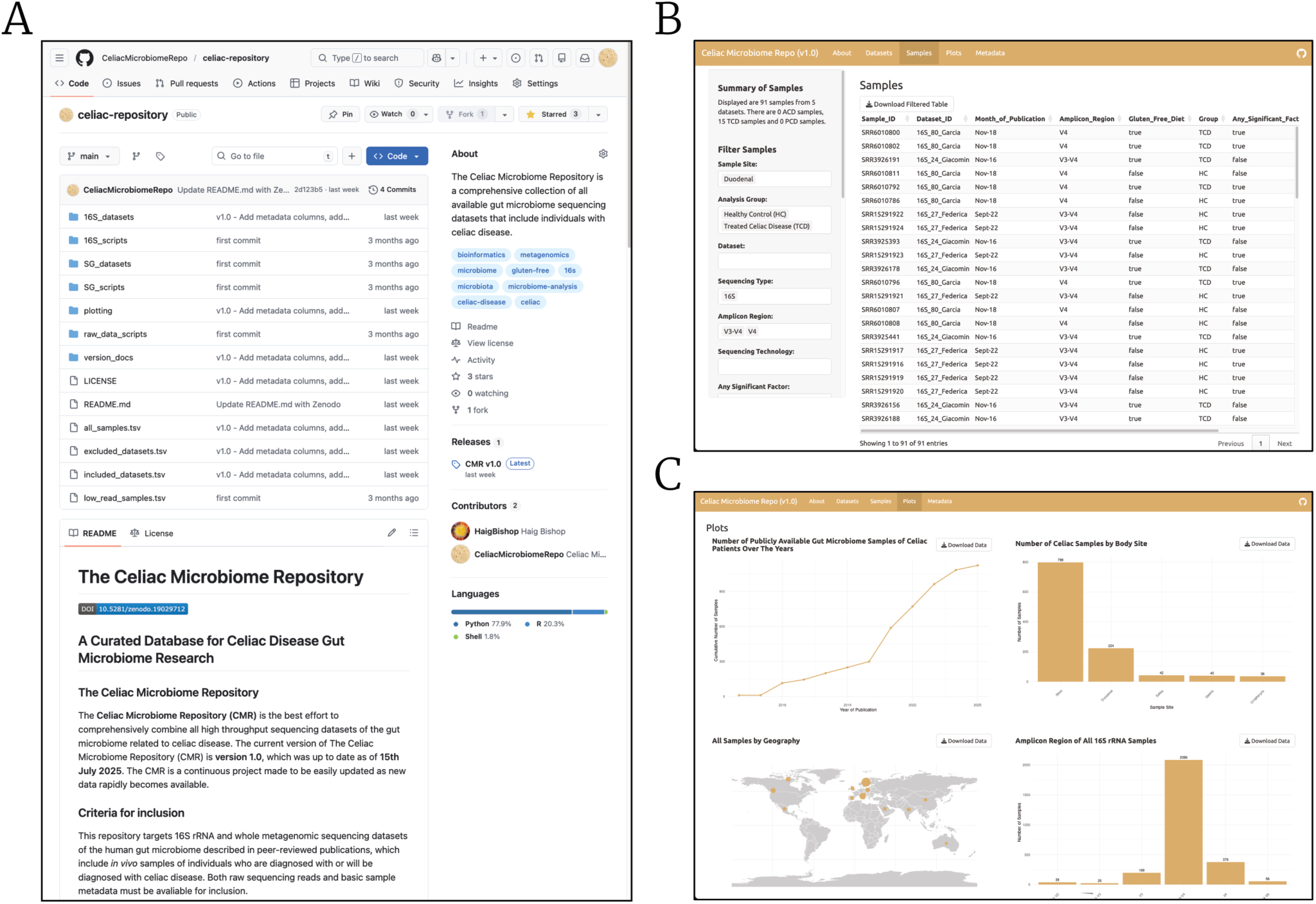
Dual-interface design of the Celiac Microbiome Repository. **A** The CMR GitHub repository homepage, housing all scripts, processed microbiome profiles, curated metadata and documentation. **B** The R Shiny web application plots tab, displaying interactive visualisations of sample and dataset distributions. **C** The R Shiny web application samples tab, providing filter controls for metadata fields and a downloadable data table of all samples. **Alt Text:** Three-panel screenshot of the Celiac Microbiome Repository dual interface: the GitHub repository homepage, the R Shiny web application plots tab showing interactive dataset and sample distribution visualisations, and the samples tab with metadata filter controls and a downloadable sample table.

At the core of the CMR is a table of all included datasets (included_datasets.tsv) and a table of all samples contained in those datasets (all_samples.tsv). These tables act as a summary of the CMR and as a starting point for the remainder of the repository’s contents. They contain a wide range of information about samples and datasets including unique identifiers, publication details, source of the data, sequencing methodology, processing details, sample demographics, and the essential metadata related to disease status and gluten-free diet. There is also a table of all datasets that were unable to be included in the CMR despite being eligible (excluded_datasets.tsv), which holds basic metadata for the publication as well as the reasons for its exclusion. The CMR includes a metadata dictionary available both as a tab on the R Shiny application and as a markdown file in the GitHub repository, clearly defining every column used across the all_samples.tsv, included_datasets.tsv and excluded_datasets.tsv files. Descriptions for these metadata fields are also provided in **Supplementary Table 4**, **Supplementary Table 5**, and **Supplementary Table 6**, respectively.

The microbiome data itself is stored in subdirectories within the repository, organised by sequencing method (*i.e.* 16S or shotgun metagenomics) and by dataset. For every 16S dataset there are various files, most importantly the unique ASV sequences (seqs.fasta), their taxonomic assignments (taxonomy.tsv) and the ASV abundance table (asv_abundances.tsv), all produced by DADA2. Similarly, for every shotgun dataset there are multiple files, most importantly a {sample_id}_profile.txt file for every sample in the dataset, which contains the sample’s microbiome community profile produced by MetaPhlAn4.

The GitHub repository also houses the Python, R and Bash scripts used for data processing. Importantly, all scripts are well-documented with inline comments and additional documentation describing the purpose of every script and how to run it. Finally, the repository contains documentation in the form of .md files, including README.md which acts as an overview of the CMR. There is also a latest_version.json file containing the version number and date of the literature search, and a repo_version_1.0.md file which describes the first release of the CMR in great detail. The purpose of this is that as new data surfaces, the CMR will be iteratively updated, and every new version should be given its own documentation.

The R Shiny web application serves as the primary discovery tool for researchers upon introduction to the repository. It allows users to immediately explore the repository’s contents without needing to write code or download large files. Users can apply filters based on metadata fields, such as body site or sequencing technology, and the application provides real-time visual summaries. For example, distribution plots are shown for amplicon regions and analysis groups, as well as plots highlighting samples with significant factors. The instant feedback allows a researcher to rapidly determine whether the CMR contains the data to power their particular investigation. Once a user has filtered the samples according to their interests, using the export functionality, they can download the curated metadata as a CSV file ready for deeper investigation.

## Discussion

### Summary of Findings

The Celiac Microbiome Repository (CMR) was developed to make the abundance of raw sequencing data from patients who developed celiac disease usable for large-scale analysis. We aggregated 28 datasets (3,245 samples) into a single, harmonised resource through literature searches and targeted author engagement. This represents a significant expansion over existing general-purpose repositories, such as GMrepo [21], which contained just four celiac-related datasets at the time of this study. This disparity highlights the significant effort required to aggregate and harmonise fragmented data, which likely explains the rarity of specialised resources such as the CMR.

A critical finding was widespread inaccessibility of both raw reads and metadata. In our search for eligible datasets, just 30 were identified via the Sequence Read Archive (SRA), whereas a more comprehensive review of the literature via Scopus was required to identify an additional 28 datasets. Of these 58 eligible datasets identified, only 20 were found to have both the necessary raw data and metadata available in the SRA or other public databases. By manually extracting metadata from publications, we made usable an additional four datasets, and direct contact with authors recovered a further four datasets. This brought the total number of usable datasets to 28, which was still less than half of the 58 datasets identified as eligible. This highlights the systemic challenge in the field where, despite the rapid data accumulation, poor adherence to data-sharing standards is preventing these samples from being used in secondary research.

By providing extensive curated metadata for every sample and dataset, the CMR transforms these isolated studies into a unified cohort suitable for high-powered meta-analyses and machine learning. Furthermore, our approach of using a standardised processing pipeline rather than pooling pre-processed data addresses the inherent batch effects of diverse bioinformatic methods. By reprocessing all 16S data through DADA2 and all shotgun data through MetaPhlAn4, we have ensured that the taxonomic profiles are methodologically consistent for cross-study comparison.

### Repository Usage and Scope

The CMR is a unified discovery and data-reuse platform that removes the need for manual literature review and metadata extraction when identifying celiac gut microbiome cohorts. By centralising fragmented resources, researchers can rapidly locate and access datasets for meta-analysis, high-powered analyses and independent exploration.

The R Shiny application enables rapid data exploration for clinicians and researchers. Users can browse and filter the 3,245 samples based on specific criteria like body site, sequencing region, or patient demographics. This allows a researcher to quickly determine, for example, how many prospective datasets target the V4 region or isolate all saliva samples from patients on a gluten-free diet. The option to export these filtered metadata tables as CSV files enables rapid cohort selection and preliminary analysis without requiring any programming expertise.

The scientific value of the CMR lies in the immediate flexibility it offers for novel research questions and analytical approaches. Because microbial profiles are standardised, researchers can move directly from cohort selection to biological inquiries and downstream analyses. This could mean using a tool like PICRUSt2 to predict functional potential in at-risk infants or running meta-analyses on specific niches like the salivary microbiome. Bioinformaticians can simply clone the GitHub repository to integrate the data directly into their own local pipelines. Additionally, the CMR offers a clear framework for benchmarking within a global context. By adopting our four-step curation workflow, other scientists can integrate their own sequencing data with the repository to see how their cohort fits into the broader celiac landscape.

Beyond traditional statistical analyses, the CMR creates the necessary foundation for machine learning. By integrating samples from 13 countries and five body sites, it encompasses geographic and methodological variance that is essential for training models that generalise. Without this consolidation, training data is limited to isolated datasets that fail to capture global diversity, making it nearly impossible to develop models that perform reliably outside of a single cohort. By providing a comprehensive, standardised collection, the research community is empowered to develop models that leverage all available data to find patterns that were previously hidden in smaller studies.

### Repository Philosophy and Cross-Field Transferability

#### Sustainable, Minimalist Architecture

Bioinformatic repositories frequently suffer from limited longevity due to lack of sustained funding or the expertise required for ongoing maintenance [58,59]. Projects like ResMicroDb utilise sophisticated stacks including MySQL, SpringBoot, and various JavaScript libraries [29]. While these offer bespoke aesthetics and advanced features such as ‘Sample Similarity Search’, they impose significant technical debt and financial cost. Their heavy reliance on third-party dependencies and advanced implementation necessitates ongoing web development expertise, and risks project abandonment if the original developer moves on. Simultaneously, the reliance on cloud providers creates a requirement for consistent funding, which introduces risk of failure if hosting fees are not met in a landscape where scientific research funding is increasingly scarce.

To mitigate these risks, the CMR employs a minimalist architecture that prioritises cost-effectiveness and simplicity. The repository is hosted through a GitHub backend and an R Shiny frontend, keeping the framework familiar and maintainable for the biology and bioinformatics community. GitHub provides free hosting for public repositories, while shinyapps.io offers a free tier for deploying Shiny applications, allowing the CMR to remain publicly accessible without maintaining bespoke infrastructure or paying for dedicated servers. This approach is sufficiently scalable for ongoing repository growth, while avoiding the complexity and technical debt of traditional web development.

#### Dual-Interface Approach

The CMR serves two types of users through a dual-interface design (GitHub backend, R Shiny frontend), addressing a common friction point in bioinformatics where data is either obscured behind programmatic barriers or trapped within rigid graphical user interfaces. By decoupling the data storage from its visualisation, the platform adapts to the technical proficiency of the user.

For clinicians and experimental biologists, the R Shiny application functions as an intuitive discovery portal. In research workflows, assessing meta-analysis feasibility requires substantial manual review and metadata extraction. The Shiny interface makes this process fast. Users can filter by body site, sequencing technology, and patient demographics to visualise what data exists, without needing R or command-line tools.

Conversely, for bioinformaticians and machine learning practitioners, GitHub provides a robust programmatic interface, avoiding per-file downloads and fragile bespoke APIs. By housing the CMR on GitHub, the entire repository, including microbiome profiles, curated metadata and processing scripts, can be cloned in a single command, facilitating immediate integration into custom computational pipelines. Ultimately, this approach democratises access to celiac microbiome data, ensuring it is as useful to both clinicians and computational researchers.

#### Adherence to FAIR Principles and Community Standards

The CMR was developed with a commitment to the FAIR (Findable, Accessible, Interoperable, and Reusable) data principles [27], ensuring the repository serves as a lasting and useful resource. Centralising celiac-related microbiome data in one GitHub repository simplifies access to fragmented datasets and fills key metadata gaps. We maximise findability by archiving each release on Zenodo with a persistent DOI and using repository tags and a detailed README. To ensure accessibility, we provide a dual-interface and release all data and code under open licences (CC BY-SA 4.0 and AGPL-3.0), removing technical and legal hurdles to data utilisation. Interoperability is achieved by providing data in standard, machine-readable formats (*e.g.*, .tsv and .fasta) and reprocessing all samples through a uniform bioinformatic pipeline to produce directly comparable taxonomic profiles. Further, we adopted GTDB taxonomy [60], a phylogenetically consistent taxonomy that is increasingly standard in microbiology, while preserving flexibility by providing original ASVs for 16S samples and both MetaPhlAn-native and GTDB-mapped profiles for shotgun samples. Finally, reusability is the primary value proposition of the CMR. Through intensive recovery, standardised processing, and metadata curation, the data can be immediately integrated into meta-analyses or used to train machine learning models with minimal additional manual work.

#### A Blueprint for Data Curation of Niche Microbiome Cohorts

While the CMR is populated with celiac-related datasets, the underlying challenges it addresses, namely dataset fragmentation and heterogeneity, are systemically shared across all microbiome research [61]. By adopting the CMR model, researchers investigating other conditions or cohorts such as Inflammatory Bowel Disease (IBD), Type 1 Diabetes, or even environmental niches such as hot spring microbial ecology, can move towards high-powered analyses and machine learning applications that leverage the full weight of available data. Because the entire system is built using open-source tools like R and GitHub, replicating the repository is a straightforward process. A researcher can fork the CMR GitHub repository and immediately have the directory structure and processing scripts required to begin their own curation. A researcher can simply replace the celiac-specific search parameters and redefine the metadata fields to suit their target cohort. For example, while the CMR prioritises gluten-free diet status, a repository for environmental samples might instead prioritise pH or geographic coordinates. Ultimately, the transferability of the CMR architecture fosters a more cooperative and data-rich future for microbiome research, by lowering the barrier to entry for large-scale meta-analyses and machine learning projects in any branch of microbiome science.

### Technical Nuances and Limitations

#### Sequencing Method Heterogeneity

While the CMR provides a harmonised framework for celiac research, several technical nuances inherent to the source data must be considered when interpreting multi-study analyses. Perhaps the most significant of these is the diversity of DNA extraction methods across the included studies. There were at least 18 unique DNA extraction kits used across the 28 datasets. It is well-documented in microbiome analysis literature that the choice of extraction kit, as well as protocol details such as the inclusion of a mechanical lysis step like bead-beating, significantly influences the observed microbial composition [62–66]. Beyond extraction methods, differences in sequencing platforms and primer choices also skew results [13,67,68]. Consequently, nearly every dataset carries a unique signature of its specific collection and processing history. While the CMR standardises the bioinformatics by re-processing all data through a uniform pipeline, it cannot retroactively eliminate these upstream biases. Consequently, the dataset variable should be treated as a random effect in CMR-based meta-analyses to control for these study-specific biases.

#### Host Contamination Challenges in Biopsy Samples

A specific technical nuance was observed in samples originating from shotgun metagenomic sequencing of duodenal biopsies collected during diagnosis. In the duodenal samples of the SG_80_Mouzan dataset [53], host DNA contamination was remarkably high, with a median of only 8.5% of reads remaining after human read removal, compared to a median of 99.9% across all other samples. This is a common challenge in biopsy-based metagenomics where human cells vastly outnumber microbial cells [69,70]. This observation highlights the unfortunate fact that the most clinically significant sampling site for celiac disease is also the least compatible with our most powerful sequencing method. Selective microbial DNA enrichment or extreme sequencing depths may offer a way to circumvent these limitations [69,70]. However, these findings serve as a warning for researchers intending to reuse this data or design similar studies on duodenal biopsies.

#### Technical Variations in 16S rRNA Gene Amplicon Targets

A significant hurdle in 16S rRNA gene meta-analysis is the lack of standardisation in the target hypervariable region during sequencing [13]. While the majority of datasets in the CMR target the V3–V4 or V4 regions, the specific primers and sequencing lengths vary. This is particularly evident in the 16S_1211_Milletich (V3–V4) and 16S_25_Francavilla (V1–V3) datasets [47,50]. In these cases, the use of single-end reads allowed for sequencing that did not span the entire target hypervariable region. When the ASVs of different datasets target different regions or incompletely overlap, taxonomic resolution can become inconsistent, since the discriminatory power of the 16S rRNA gene is unevenly distributed across its nine hypervariable regions [71]. This can also create significant challenges when attempting to truncate ASVs across datasets for integration, since the actual sequence coverage may deviate from the target hypervariable regions. By using the extract16s tool to align all ASVs to a reference *E. coli* sequence, we precisely mapped these discrepancies, providing researchers with the transparency required to avoid potential pitfalls when conducting multi-study analyses.

While full-length 16S rRNA gene sequencing was not utilised by any studies in version 1.0 of the CMR, it is an increasingly viable approach that achieves higher taxonomic resolution compared to short-read 16S rRNA gene amplicon sequencing [72,73]. We recommend that future 16S-based studies adopt full-length sequencing where possible to mitigate the resolution limits and biases associated with relying on fewer regions [13,14]. If full-length sequencing is not feasible, the V3–V4 hypervariable region remains the preferred target, since it is well established in gut microbiome research, provides relatively high discriminatory power, and maintains broad phylogenetic coverage [13,74,75]. Additionally, as the most common region in current CMR datasets, it ensures maximum consistency with existing celiac microbiome data.

#### Blind Spots in the Celiac Microbiome Landscape

While the CMR represents the most comprehensive collection of celiac microbiome data to date, it also exposes some “blind spots” in the research landscape. Geographically, the data is skewed toward Western, industrialised nations. Of the 28 included datasets, the vast majority originate from Europe and North America, leaving a void of data in South America, Africa, and parts of Asia where celiac disease is now recognised as a significant public health problem that is no longer restricted to those of European ancestry [5]. This disparity reflects the geographic bias towards highly developed countries in the field, with over 71% of available public microbiome samples originating from Europe and North America [76]. Because geography has such a significant impact on the microbiome [77], the repository’s current lack of global representation inevitably limits the generalisability of meta-analyses or machine learning models trained on the CMR to lesser-represented geographies and ethnicities.

The data imbalance extends beyond geography to the fundamental sequencing strategies and study designs. Only five of the 28 datasets employ shotgun metagenomics, while three of the ten most recent additions use this method. Clearly the repository is still dominated by 16S rRNA gene amplicon data, restricting the functional and strain-level resolution available across the collective samples. Similarly, the scarcity of prospective studies remains a bottleneck. With only four of these datasets and 136 prospective celiac samples, the repository is primarily composed of cross-sectional snapshots that cannot distinguish between microbial causes and effects of celiac disease. Determining this directionality is vital for moving beyond describing the diseased state toward identifying early microbial drivers of celiac pathogenesis. Notably, at the intersection of shotgun metagenomic sequencing and prospective studies, there is just one dataset, underscoring the shortage of precise longitudinal evidence required to map specific strains and functional pathways to the onset of celiac disease.

Based on the current dataset composition, the research community would most benefit from studies that expand into underrepresented geographies and employ shotgun metagenomic sequencing. In particular, shotgun sequencing of prospective cohorts and duodenal biopsies would provide the most novel mechanistic insights into the gut microbiome’s role in celiac disease, provided that researchers carefully manage the high levels of host DNA contamination common in biopsy samples. Beyond diversity, the CMR remains open to all data types, since greater overall volume will inevitably strengthen the collective statistical and predictive power of future meta-analyses and machine learning models.

#### Metadata Standardisation Challenges

Despite extensive curation effort, there remain several limitations of the resulting metadata that are important to note. Basic demographic variables such as age and sex were frequently omitted from public records, preventing their use in downstream analyses. Furthermore, critical clinical markers including HLA haplotypes and Marsh classification scores, which grade the extent of villous atrophy in duodenal biopsies, were so infrequently reported that they were not included as standard fields. This lack of granular clinical data makes it impossible to distinguish between patients with varying degrees of intestinal damage or HLA haplotype, obscuring any specific microbial signatures associated with disease severity or genetics. Future studies should, where possible, prioritise the collection and reporting of core demographic and clinical variables, specifically age, sex, celiac diagnosis, gluten-free diet status, HLA haplotypes, and Marsh scores. To ensure availability, we recommend that researchers provide this metadata when depositing raw sequencing reads into public repositories like the SRA.

Beyond missing data, the lack of universal definitions for key variables such as celiac diagnosis and gluten-free diet presented a challenge. The criteria for a celiac disease diagnosis vary over time and across different regions [78], so studies in the CMR are liable to use different diagnostic rules. However, the inherent constraints of the available data necessitated a reliance upon the diagnoses provided by the original study, while acknowledging the potential diagnostic variation. A similar challenge exists for gluten-free diet status. While adherence is rarely a binary state, a lack of data regarding diet duration or strictness forced us to adopt a simplified binary view. This lack of standardisation of metadata definitions means the CMR aggregates a wide spectrum of clinical phenotypes under broad labels. While this provides unprecedented scale, users should interpret results as representing an aggregate biological signal rather than precise clinical categories.

### Future Expansion and Maintenance

The CMR was created to be a living repository rather than a static archive. We commit to performing annual updates that mirror the outlined four-step workflow to identify and incorporate new datasets for as long as the repository is actively used. To support reproducibility, each update will be archived on Zenodo and receive a unique DOI, which researchers can use to reliably cite the specific CMR version used. While our curation process for version 1.0 revealed a large number of inaccessible datasets, we will maintain the capacity to include these in future versions. We encourage authors of the excluded datasets to reach out via email to share their data, helping expand the repository’s global coverage.

Beyond routine updates, the nature of GitHub as the primary platform makes collaboration and expansion straightforward. We welcome contributions from the wider research community, whether through identifying new data or proposing extensions to the repository’s scope. Future iterations could integrate functional profiling tools such as PICRUSt2 or HUMAnN3 to provide insights into metabolic pathways and gene abundances alongside existing taxonomic data. Also, as sequencing technologies evolve, the CMR can adapt accordingly, for instance by incorporating additional bioinformatic tools to better support long-read sequencing. Further, future updates could expand the collection of celiac-specific metadata to higher-fidelity records of Marsh scores and HLA haplotypes, or broaden eligibility criteria to encompass conditions such as non-celiac gluten sensitivity. By providing a transparent and modular framework, we invite other researchers to help the CMR evolve alongside the rapidly changing landscape of microbiome science.

## Conclusion

The Celiac Microbiome Repository (CMR) represents a fundamental shift in how the research community can interact with celiac disease sequencing data. Prior to this work, the celiac microbiome landscape was fragmented and valuable samples remained dormant in public archives. By harmonising 28 datasets and 3,245 samples into a single, open-access resource, we have transformed these isolated snapshots into a unified cohort.

Our dual-interface architecture ensures that this data is not just findable, but truly functional. The R Shiny application offers a window into the global state of celiac microbiome research without the need for computational expertise, while the GitHub backend provides a foundation for future meta-analyses and machine learning models. By lowering the technical and administrative barriers to data reuse, we provide the community with the tools to move beyond small-scale studies and toward the global, high-powered discoveries required to fully understand the microbial drivers of celiac disease. We invite the research community to explore and build on this repository as we work toward a more collaborative, data-rich future for microbiome science.

## Author Contributions

Bishop H.V. conceived the study, conducted the data retrieval, curation and processing, performed the analyses and wrote the manuscript. Prendergast P.J. assisted in the data retrieval, curation and processing. Herbold C.W., Dobson R.C.J., and Ogilvie O.J. supervised the work, provided critical input on the study design, and revised the manuscript. Dobson R.C.J. and Ogilvie O.J. secured research funding.

## Code and Data Availability

The data retrieval and processing pipeline scripts along with all microbiome profiles and curated metadata are available on the CMR GitHub repository (https://github.com/CeliacMicrobiomeRepo/celiac-repository). To ensure long-term persistence and reproducibility, every version of the repository is archived on Zenodo (https://doi.org/10.5281/zenodo.19029712). The source code for the R Shiny application is also hosted on GitHub (https://github.com/CeliacMicrobiomeRepo/celiac-webapp). All scripts housed in the CMR GitHub repository, and the R Shiny source code are provided under the AGPL-3.0 licence, while curated metadata and processed microbiome profiles are provided under the CC BY-SA 4.0 licence.

